# Modular Nanobody Conjugates with Controlled Topology Using Genetically Encoded Non-canonical Amino Acids

**DOI:** 10.1101/2025.11.27.691038

**Authors:** Roman Adomanis, Nathan Phan, Grant Walter, Blaise R. Kimmel

## Abstract

Bispecific antibody-based derivatives are traditionally generated by fusing short, flexible peptides to two variable heavy and light chain pairs that recognize distinct antigens. However, this method limits domain joining to the amino (N) or carboxyl (C) termini, thereby restricting our ability to study the role of domain orientation in key properties of bispecific molecules, such as affinity and specificity. Here, we present an adaptable, plug-and-play application of genetic code expansion technology for the rapid, modular creation of bispecific nanobody conjugates from non-canonical amino acid (ncAA)- integrated nanobody domains, offering precise control over domain topology. We implement this strategy using a single-plasmid genetic code expansion system and computation-guided ncAA site selection to preserve nanobody expression and target binding. We demonstrate the effective incorporation and crosslinking of azide- and tetrazine-modified lysine and phenylalanine, respectively, at four engineered positions within an anti-PD-L1 nanobody and two positions within an anti-CTLA-4 nanobody in a panel of immune cell- and cancer-cell-binding nanobodies. Using this approach, we demonstrate the modular synthesis of a library of bispecific nanobodies, comprising four PD-L1xCD16 immune synapse engagers and eight PD-L1xCTLA-4 immune checkpoint bispecifics. This work enables a new dimension of control over protein crosslinking for rapidly constructing immune cell engagers for dual immune checkpoint blockade therapy and to bridge the synapse between immune cells and cancer cells.

## Introduction

The recombinant expression of fusion proteins is limited to the use of flexible peptide linkers to genetically fuse multiple protein domains^1^. While this approach offers a stable covalent linkage, a sufficiently long and flexible linker must separate the protein domains, and expression is limited to canonical N-to-C-terminal rules, producing only fusions that differ only in the linear arrangement of domains. Therefore, in two-component systems (e.g., bispecific scFvs), only two isoforms (A-B and B-A) are possible, limiting the therapeutic impact of domain topology for these largely understudied molecules. Recent work in the protein immunotherapy space has highlighted the importance of geometric control over targeting domains, demonstrating orientation-dependent improvements in antigen affinity and virus neutralization for a nanobody surface-functionalized nanoparticle and a tri-specific nanobody conjugate, respectively^2,3^.

To achieve this geometric control, individual antigen-targeting domains with reactive handles can be expressed independently and then covalently crosslinked using polymeric linkers^4,5^. However, to ensure high product homogeneity, these handles must be orthogonal to each other, and ideally, react rapidly under physiological conditions. Among strategies for chemically linking recombinant proteins, one of the most promising approaches has been spontaneous, orthogonal click-chemistry strategies, including strain-promoted azide-alkyne cycloaddition (SPAAC) and inverse electron-demand Diels-Alder cycloaddition (IEDDA) reactions^6,7^. IEDDA click reactions are highly selective and exhibit rapid conjugation kinetics, with reported second-order rate constants on the order of 10^^6^ M^-1^ s^-1^, which have been shown to improve the crosslinking of protein domains and to offer precise labeling of chemical agents (e.g., chemotherapy, immunotherapy) for targeted protein delivery^8–10^. However, due to limitations of site-specific integration of these functional click-chemistry handles into proteins at the N’ and C’ terminals, the resulting bispecific products resemble genetically fused isoforms, which continues to limit the optimal geometric configurations of the protein domains.

Here, we present a modular protein bioconjugation strategy that improves the geometric configuration of protein domains, thereby optimizing protein function upon tethering, by integrating non-canonical amino acids (ncAAs)^11^. These ncAAs offer distinct chemistries from the canonical 20 amino acids within the protein domains, with recent work showing the design and integration of residues capable of participating in both SPAAC and IEDDA click chemistry reactions. Inspired by this work that motivates the integration of these ncAAs into proteins for site-specific chemical modification of proteins at any site – rather than just the N’ or C’ terminal – we engineered a platform technology that showcases the rapid construction of bispecific proteins after recombinant expression of each subunit domain using a single plasmid system that integrates the necessary machinery (i.e., aminoacyl-tRNA synthetase/suppressor tRNA pairs; aaRS/tRNA) required to achieve genetic code expansion (GCE)^12^. We used molecular protein design strategies (e.g., Rosetta)^13^ to identify optimal sites for ncAA integration into a recently reported single-domain antibody fragment (i.e., nanobody) that binds PD-L1 receptors (nPD-L1) on cancer cells and myeloid cells. With the key advantages afforded via site-specific control over ncAA incorporation, we integrate this GCE technology into a modular platform using the optimal four nPD-L1 candidates to generate four geometrically distinct fusions with an anti-CD16 (nCD16) natural killer (NK) cell binder, and eight geometrically distinct fusions to an anti-CTLA-4 nanobody (nCTLA-4) for targeting lymphocytes (T cells)^14,15^. We demonstrate the construction of bispecific nanobodies with unique topologies that were crosslinked using a commercially available heterobifunctional linker to generate fusions with precision over the sites of chemical crosslinking, affording a homogeneous product. Collectively, our technology highlights the potential of this platform for high-throughput drug discovery applications by demonstrating high reaction efficiency, product homogeneity, and rapid purification of our fusions, positioning our approach for rapid bispecific protein synthesis for *in vitro* screening to inform critical decisions on domain orientation for new bispecific immunotherapeutics^16^.

## Results

### Construction of a Single Plasmid Genetic Code Expansion System

This research builds on the previous work of the Mehl group to engineer an orthogonal aaRS/tRNA pair for the robust incorporation of several meta-substituted 1,2,4,5-tetrazine-phenylalanine derivatives in both *E. coli* and eukaryotic cells^17,18^. Here, we focus on the application of the 6-butyl-1,2,4,5-tetrazine-containing phenylalanine derivative (Tet3.0 Bu; Tet3.0) **(Figure 1B)** due to its high incorporation efficiency and rapid reaction kinetics with strained alkynes^19^. To control the homogeneity of our bispecific conjugate products, we turned towards the azide-containing pyrrolysine derivative N^ε^-((2azidoethoxy)carbonyl)-L-lysine (AzK) **(Figure 1B)**, a substrate of the well-characterized wild-type *Methanosarcina mazei* pyrrolysyl-tRNA synthetase/tRNA pair (*Mm*PylRS/*Mm*tRNA_CUA_^Pyl^). AzK serves as an orthogonal reaction site to Tet3.0, demonstrating rapid and selective cycloaddition with DBCO. Low ncAA incorporation efficiency, resulting in yields 30-80% lower than wild-type expression, remains a key barrier to the widespread application of GCE technology^17,20^. While several groups have attempted to address this challenge using approaches such as orthogonal ribosome systems or genome-wide knockout of release factor 1 (RF1) in expression hosts, we chose to focus on recent work by the Wiltschi group to reduce the metabolic burden associated with commonly used multi-plasmid systems for GCE^18^. The Wiltschi group demonstrated that the colocalization of the *Mm*PylRS, *Mm*tRNA_CUA_^Pyl^, and protein of interest (POI) genes, all under control of the T7_*lacO*_ promoter (T7p) and terminator (T7t), resulted in a significant elevation of EGFP yield in the presence of AzK. Therefore, we constructed pRESS, a <u>p</u>rokaryotic <u>R</u>ecombinant <u>E</u>xpression and <u>S</u>uppression <u>S</u>ystem, based on the established pET28b(+) plasmid **(Figure 1C)**. Like the work of Galindo Casas *et al*.^21^, pRESS contains genes for *Mm*PylRS, *Mm*tRNA_CUA_^Pyl^, and the POI, all under control of T7p/T7t and regulated by the *lacI* gene. Notably, to ensure proper 5’ and 3’ processing of the *Mm*tRNA_CUA_^Pyl^ precursor processing, the *proK* promoter and terminator were retained in the gene cassette and flanked by T7p/T7t, rather than replacing them. Importantly, pRESS contains orthogonal golden gate cut sites, *BsaI* and *PaqCI*, flanking the POI and *Mm*PylRS sequences, providing a modular platform for incorporating a wide range of ncAAs by inserting mutant *Mm*PylRS genes **(Figure 1C)**.

**Figure 1:**
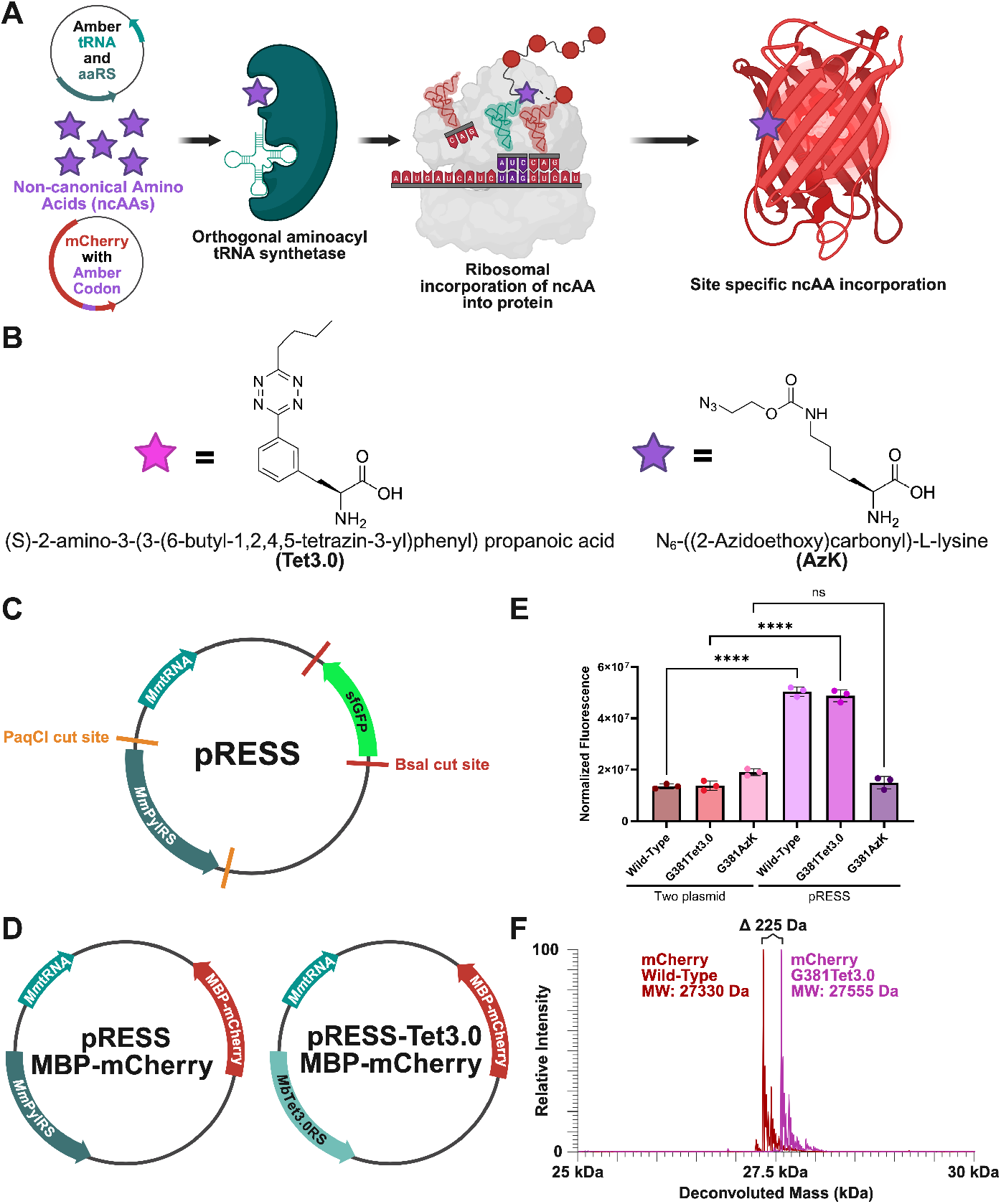
Construction of a single plasmid GCE system. **(A)** Traditional two-plasmid systems for GCE in which the aaRS/tRNA pair is expressed on a separate plasmid from the gene of interest. (**B)** Chemical structures of the two ncAAs used in this work, referred to as Tet3.0 and AzK. (**C)** The pRESS plasmid is used to express proteins of interest with site-specific incorporation of ncAAs. (**D)** The reporter pRESS MBP-mCherry and pRESS-Tet3.0 MBP-mCherry plasmids after construction using the *BsaI* or *PaqCI* sites for Golden Gate assembly. (**E)** MFI data of cultures grown in the presence of 1 mM of Tet3.0 or 2 mM AzK 72 hours post-induction, demonstrating improved MBP-mCherry^G381TAG^ expression using pRESS. **F)** ESI-MS data of MBP-mCherry^WT^ and MBP-mCherry^G381Tet3.0^ demonstrating the ncAA incorporation efficiency of pRESS.

To demonstrate the modularity and reduced metabolic burden of pRESS, we created pRESS-Tet3.0, which contains the *Methanosarcina barkeri-*derived pyrrolysyl-tRNA synthetase (*Mb*Tet3.0RS) developed by the Mehl group^10^. A his-tagged maltose-binding protein (MBP) domain, followed by a tobacco etch virus (TEV) protease cleavage site and a glycine-serine linker (G_4_S), was fused to the N-terminus of mCherry to create the fluorescent reporter MBP-mCherry. Wild-type (MBP-mCherry^WT^) and mutant MBP-mCherry (MBP-mCherry^G381TAG^), harboring the TAG codon at position G381, which is placed three amino acids upstream of the mCherry domain, were inserted into pRESS and pRESS-Tet3.0 and transformed into the *E. coli* expression strain BL21(DE3) **(Figure 1D)**. In parallel, MBP-mCherry^WT^ and MBP-mCherry^G381TAG^ were inserted into the parental pET28b(+) plasmid and co-transformed into BL21(DE3) cells along with pUltraI-Tet3.0 [TAG] or pUltraI-*Mm*Pyl [TAG], derivatives of the pUltraI-Tet3.0 [TAA] plasmid developed by the Mehl and Schultz groups, which we modified to recognize the TAG rather than the TAA codon and/or contain genes for *Mm*PylRS and *Mm*tRNA_CUA_^Pyl^ in the case of pUltraI-*Mm*Pyl^10,19^. The fluorescence intensity of each culture, normalized to OD600, measured 72 hours post-induction, shows a significant improvement in expression of both wild-type and Tet3.0-containing mCherry, with no improvement in expression of Azk **(Figure 1E)**. The high suppression efficiency and site-selectivity of our pRESS system were confirmed using Electrospray ionization mass spectrometry (ESI-MS) analysis of MBP-mCherry^WT^ and MBP-mCherry^G381Tet3.0^, following purification and removal of the MBP domains, as evidenced by the expected mass difference of 225 Da between mCherry^WT^ and mCherry^G381Tet3.0^ **(Figure 1F)**.

### Rational Design of ncAA-Containing Nanobody Domains

Following pRESS validation, we sought to develop a method for rationally selecting sites for ncAA incorporation within our nanobody domains to build a small panel of unique bispecific molecules without loss of function. To reduce the dimensionality of our problem and establish design guidelines, we began our investigation with a single nanobody (anti-PD-L1), based on our previously reported sequence^4^. To help identify prime candidate sites for non-terminal ncAA insertion within nPD-L1, we performed *in silico* modeling of potential mutant nPD-L1 structures containing AzK using the PyRosetta software package. Several residues within the framework and complementarity-determining regions (CDRs) of nPD-L1 were mutated to the cycloaddition product between AzK and DBCO, and the resulting mutants were evaluated based on two major criteria: the change in binding free energy to the target antigen (binding ΔΔG) and the change in folding free energy (folding ΔΔG). Although the output of these metrics cannot be directly translated into kcal/mol without iterative experimental validation, they can provide intuitive guidance on ncAA site selection based on consistent principles of thermodynamic favorability and the structure-function relationship dogma of biology. Among the candidate sites identified, we chose to mutate R45 and K75 due to their minimal disruption of binding ΔΔG, structural similarity to AzK, and their distal positioning relative to each other and the CDRs **(Figure 2A-C)**. In addition to these framework sites, the first and last amino acids of the canonical nPD-L1 sequence were replaced with the TAG codon to mimic the geometric orientation of traditional genetic fusions and produce nPD-L1^Q1Azk^ and nPD-L1^S116Azk^ **(Figure 2A, Figure 2C)**.

**Figure 2:**
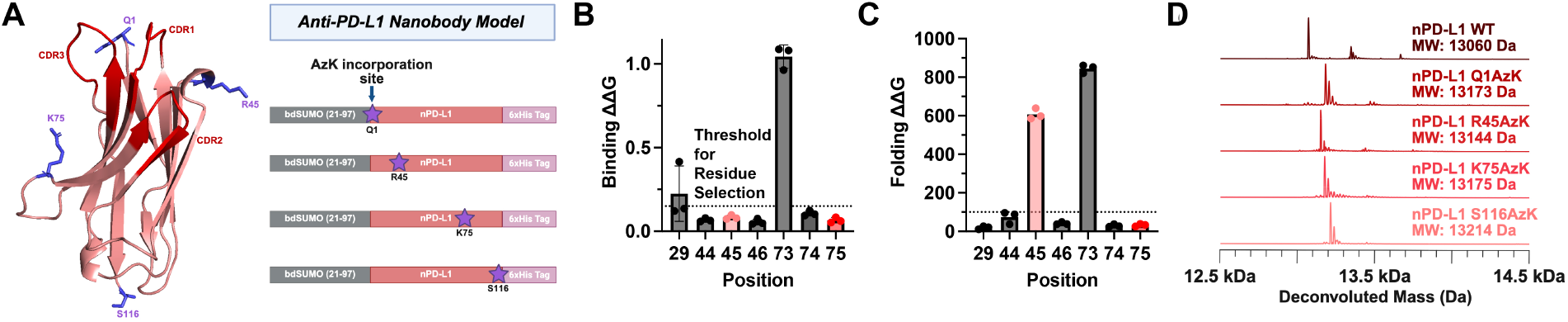
Mutation of nanobody framework residues. **(A)** Structure of nPD-L1 modeled in Rosetta with sites for ncAA incorporation highlighted, with the favorability of these sites for mutation quantified using scores for (**B)** binding ΔΔG and (**C)** folding ΔΔG. (**D)** ESI-MS data validating the ncAA incorporation for nPD-L1Q1^Azk^, nPD-L1S116Azk, nPD-L1^R45AzK^, and nPD-L1^K75AzK^. For a full list of expected MW of proteins in this study (**Table S1**).

A cleavable N-terminus *<u>B</u>rachypodium <u>d</u>istachyon*-derived <u>S</u>mall <u>U</u>biquitin-like <u>MO</u>xdifier (bdSUMO) domain and C-terminus 6xHis tag were added to address challenges in N- or C-terminally placed ncAAs **(Figure 2A)**. Insertion of a stop codon following the canonical ATG start codon shifts translation to a downstream initiation site, making N-terminal ncAA placement non-trivial^22,23^. Conversely, pre-mature truncation products for C-terminus ncAA sites would be nearly identical in size and chemical composition to the full-length ncAA-containing target protein, making their separation challenging. By including the ncAA incorporation site before the 6xHis tag, only full-length products can be easily purified via immobilized-metal affinity chromatography. ESI-MS analysis of these samples revealed high homogeneity and ncAA incorporation, consistent with the MBP-mCherry reporter results **(Figure 2D)**. It is worth noting that yields for nPD-L1^Q1AzK^ and nPD-L1^R45AzK^ were consistently higher, about 2.8 mg per liter of culture, compared to those for nPD-L1^K75AzK^ and nPD-L1^S116AzK^, which typically ranged from 0.5 to 0.6 mg per liter of culture, indicating the potential impact of folding ΔΔG scores on experimental yields.

### Modular Assembly of Bispecific Isoforms Using Nanobody Building Blocks

Following the computational design of optimal integration sites within the model nPD-L1 nanobody, we inserted pRESS constructs encoding the previously described nPD-L1 mutants and a single mutant of a previously described anti-CD16 (nCD16) nanobody for insertion of AzK and Tet3.0, respectively, to create a small panel of unique bispecific molecules^24^. For the nCD16 mutant, we chose to replace only the first amino acid, creating nCD16^E1Tet3.0^, as a proof-of-concept for our technology to produce bispecific molecules. Alone, nPD-L1 and nCD16 provide discrete molecular functions for connecting synapses between cells, where nPD-L1 binds to PD-L1 expressed receptors in cancer cells and myeloid cells (e.g., dendritic cells), and where nCD16 binding to CD16 receptors on natural killer (NK) cells offers a strategy to redirect NK cells and activate NK cells during the binding event **(Figure 3B)**. As with the nPD-L1 mutants, ESI-MS confirmed selective site incorporation of Tet3.0 into nCD16^E1Tet3.0^ with a yield of approximately 3.3 mg per liter of culture **(Figure 3B)**.

**Figure 3:**
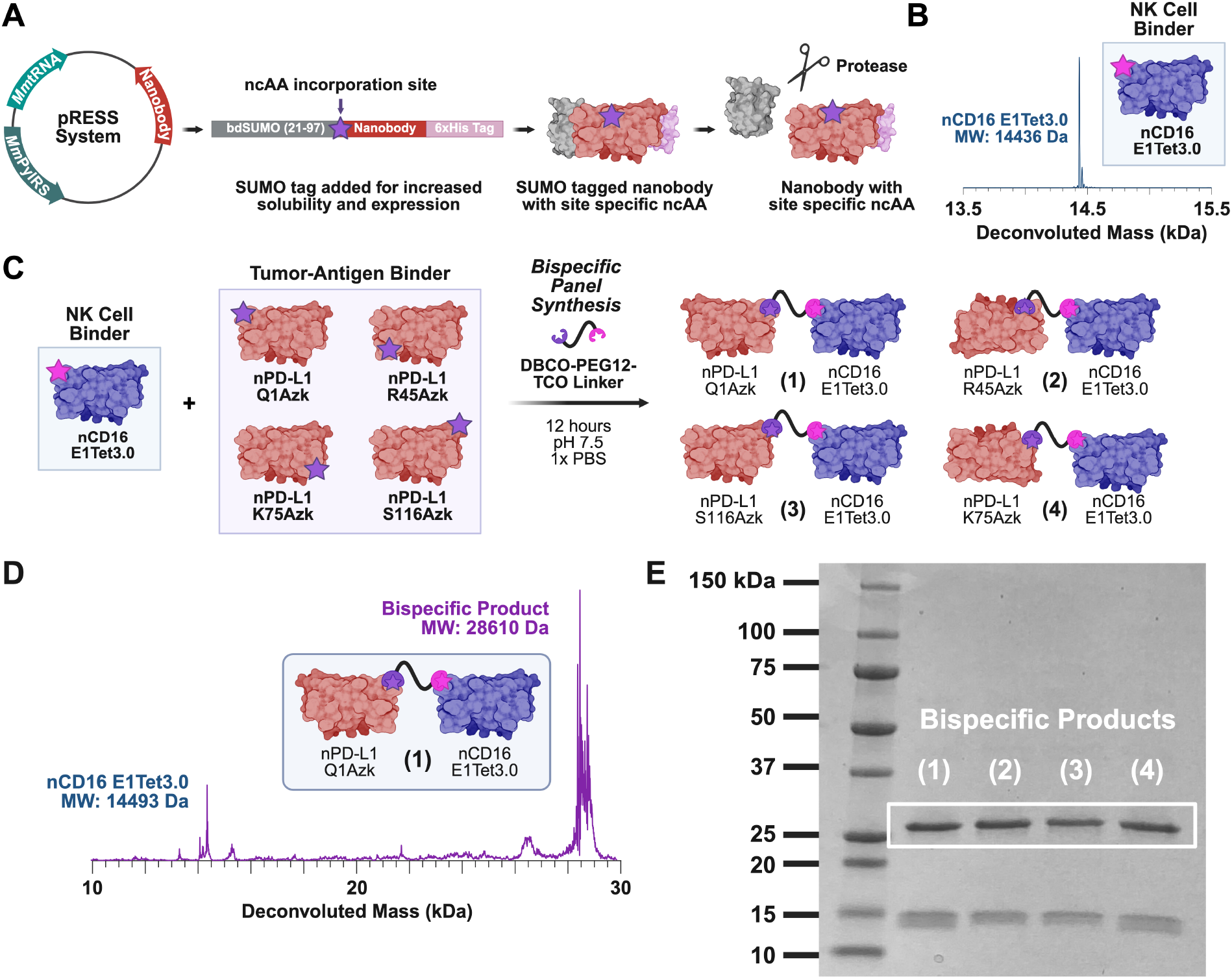
Crosslinking and characterization of anti-PD-L1xCD16 fusions using nanobody building blocks. **(A)** Expression cassette featuring an N-terminal bdSUMO domain and a C-terminal 6xHis used for all nanobody mutants in this study. (**B)** ESI-MS data validating ncAA incorporation for nCD16^E1Tet3.0^ and **(C)** the crosslinking scheme using the commercially available TCO-PEG12-DBCO linker at physiological conditions to create geometrically unique fusions **(1)-(4)**. (**D)** Representative ESI-MS data of unpurified fusion **(1)**. (**E)** SDS-PAGE showing unpurified fusions **(1)-(4)**

To investigate our capacity to make functional ncAA-based bispecific conjugates, we began by creating a panel of all possible permutations using the mutant nPD-L1 and nCD16 domains. Each nanobody combination was reacted with a commercially available heterobifunctional TCO-PEG12-DBCO linker in a 1:1:1 molar ratio and incubated overnight at room temperature under physiological conditions **(Figure 3B-C)**. The resulting isoforms **(1)-(4)** were produced with high efficiency and product homogeneity, as characterized by SDS PAGE and ESI-MS, and were easily separated from the remaining starting material using size exclusion chromatography **(Figure 3D-E)**.

Based on the results of our anti-PD-L1xCD16 bispecific assemblies, we sought to highlight the modularity of our platform by updating our panel to include an alternative immune cell engager to nCD16. We chose a previously described anti-CTLA-4 nanobody (nCTLA-4) due to the reported anti-tumor effects of the nanobody, and cross reactivity to human and murine CTLA-4^25^. Alone, nPD-L1 and nCTLA-4 act as antagonists of their respective immune checkpoints, thereby improving T-cell activation and function in immunosuppressive environments, such as the tumor microenvironment. As a fusion, this molecule has the potential to function as both a dual immune checkpoint inhibitor and a redirector of CTLA-4-positive T cells toward PD-L1-expressing cells, thereby increasing immunological activity in suppressive tumors. As with nPD-L1, the first and last amino acids of the canonical nCTLA-4 sequence were replaced with the TAG codon to produce nCTLA-4^Q1Tet3.0^ and nCTLA-4^S123Tet3.0^ **(Figure 4A-B)**. ESI-MS analysis of these mutants revealed high homogeneity and ncAA incorporation, consistent with the results obtained for all previous proteins **(Figure 4B)**. Yields for nCTLA-4^Q1Tet3.0^ and nCTLA-4^S123Tet3.0^ were also relatively low, at 0.75 to 0.5 mg per liter of culture, respectively. Like with our anti-PD-L1xCD16 bispecific panel, all possible permutations of mutant nPD-L1 and nCTLA-4 were reacted with a commercially available heterobifunctional TCO-PEG12-DBCO linker in a 1:1:1 molar ratio and incubated overnight at room temperature under physiological conditions **(Figure 4A)**. The resulting isoforms **(5)-(12) (Figure 4C-D)** were produced with high efficiency and product homogeneity, as characterized by ESI-MS.

**Figure 4:**
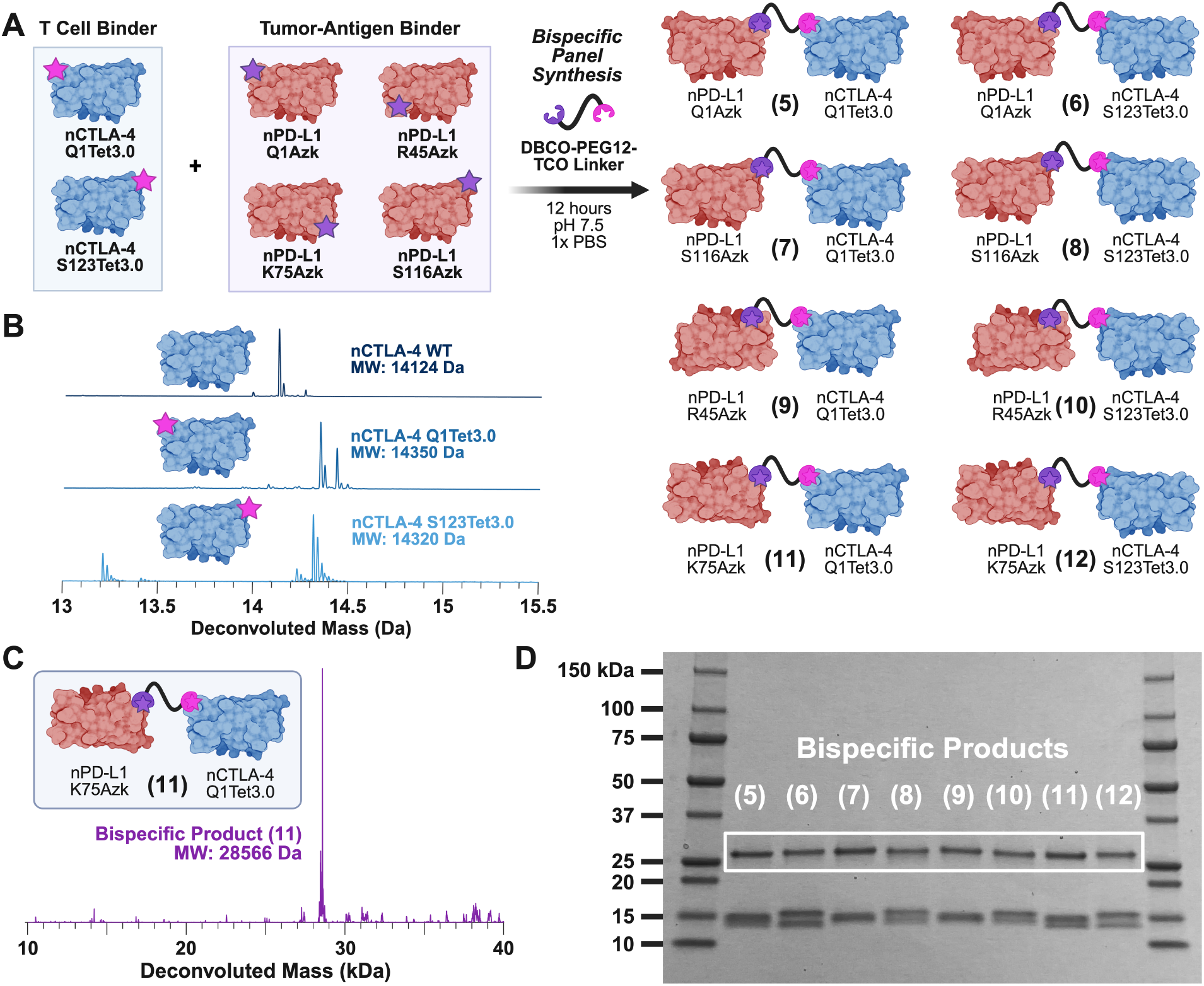
Crosslinking and characterization of anti-PD-L1xCTLA-4 fusions using nanobody building blocks. **(A)** Crosslinking of nPD-L1^Q1Azk^, nPD-L1^S116Azk^, nPD-L1^R45Azk^, nPD-L1^K75Azk^, nCTLA-4^Q1Tet3.0^, and nCTLA-4^S123Tet3.0^ using the commercially available TCO-PEG12-DBCO linker at physiological conditions to create geometrically unique fusions **(5)-(12)**. (**B)** ESI-MS data validating ncAA incorporation nCTLA-4^Q1Tet3.0^, and nCTLA-4^S123Tet3.0^. (**C)** Representative ESI-MS data of purified fusion **(11)**. (**D)** SDS-PAGE showing unpurified fusions **(5)-(12)**

## Discussion

Within the past decade, bispecific antibodies (BsAbs) and antibody fragments have become increasingly relevant therapeutic molecules in immunotherapy, regulating cell synapse formation by suppressing existing connections (i.e., checkpoint blockade) and improving immune cell redirection (i.e., T cell engagers)^26,27^. In particular, since the approval of blinatumomab in 2014, the FDA has approved 10 additional bispecific antibody-based therapies for the treatment of cancer, hematologic, and ocular diseases^28^. For solid tumors, the dual administration of antibodies targeting programmed death 1 (PD-1)/programmed death ligand 1 (PD-L1) immune checkpoints, as well as cytotoxic T lymphocyte-associated protein 4 (CTLA-4), has shown potent efficacy and safety in several cancer types^25^. However, this dual treatment regimen is more expensive and associated with a higher incidence of adverse events than single-agent therapies, making PD-1/PD-L1 and CTLA-4 increasingly attractive targets for bispecific therapies. Recently, a novel anti-PD-L1 and CTLA-4 bispecific antibody, KN046, was well tolerated and demonstrated promising antitumor efficacy in patients with advanced solid tumors in a Phase I clinical trial^29^. Another emerging topic in the field of BsAbs has been the development of novel, bispecific, heavy-chain-only single-domain antibodies, or nanobodies. A significant driver of this trend is the structural simplicity and high physicochemical stability of nanobodies, which enable efficient production in microbial hosts^30,31^. This results in significantly lower manufacturing costs, complexity, and time compared to standard mAbs and antibody fragment formats, such as single-chain variable fragments (scFvs)^32,33^. Our work aligns with this trajectory in the field by using nanobody building blocks as themolecular basis for bispecifics assembled via ncAA-enabled conjugation chemistry^16^.

In this work, we demonstrate the adaptation of GCE technology to a high-throughput platform for the creation of novel bispecific therapeutics. We highlight the modularity of our system by rapidly generating unique isoforms of nanobody-based PD-L1xCD16 and PD-L1xCTLA-4 bispecifics to mimic immune cell engagers and immune checkpoint blockade therapies, without requiring a new plasmid for each combination, and by allowing combinations to form these conjugates with uniquely defined geometric orientations of each domain. The use of ncAA-based conjugation enables rapid assessment of key physicochemical properties, such as conjugate linker length, flexibility, and individual domain orientation, on therapeutic outcomes without the need to create new expression plasmids. In particular, the PD-L1xCTLA-4 isoform panel, generated from a standard set of nanobody building blocks, provides a direct way to compare how bispecific domain geometry correlates with dual-target binding, cell-bridging, and cytolysis readouts *in vitro*. Collectively, our study establishes a foundation for future *in vitro* and preclinical development of modular bispecific protein conjugates, in which newly developed nanobody candidates can be engineered, conjugated, and screened to investigate the roles of domain avidity, affinity, and antigen recognition in cancer immunotherapy^34^.

## Methods and Materials

### pET28b Plasmids Construction

A pET28b+ plasmid, reported from recent work^35,36^, contains a constitutively expressed sfGFP gene flanked by *BsaI* cut sites. A T7 promoter-lac operator and a T7 terminator are located upstream and downstream, respectively, of these *BsaI* sites for replacement of the sfGFP gene with genes of interest under the control of the T7 promoter-lac operator. To insert genes for MBP-mCherry^WT^ and MBP-mCherry^G381TAG^ into the pET28b backbone, gBlocks containing terminal *BsaI* cut sites and an N-terminal 6xHis-tag affinity tag for purification via immobilized metal affinity chromatography (IMAC) for each protein were synthesized (IDT) and ligated into the vector via a *BsaI* Golden Gate reaction from the NEBridge® Golden Gate Assembly Kit protocol (NEB #E1601L). Constructs were then transformed into chemically competent DH10β E. coli (NEB #C3019H), then plated at 37°C on LB-Kan (50 mg/mL) selection media for 12-16 hours. Colonies were inoculated in LB-Kan (50 mg/mL) and grown at 37°C (250 rpm) for 12-16 hours, and then miniprepped using the Qiagen QIAprep Spin Miniprep Kit (Qiagen #27104). The sequence of the constructs was then confirmed using Genewiz Plasmid-EZ whole plasmid sequencing.

### pUltra Plasmid Construction

The pUltraI-Tet3.0[TAA] plasmid was a gift from Ryan Mehl (Addgene plasmid #164580). This plasmid originally contains a single copy of a Methanosarcina barkeri-derived pyrrolysyl aminoacyl tRNA synthetase (Tet3.0RS) that demonstrates polyspecificity to several meta-substituted 1,2,4,5-tetrazine-phenylalanine derivatives under control of the tacI promoter and a single copy of the cognate tRNA recognizing the ochre stop codon (tRNA^UUA^) under the control of the proK promoter. To repurpose this plasmid for the incorporation of our target ncAAs, we replaced both the aminoacyl tRNA synthetase and tRNA genes (pUltraI-*Mm*Pyl [TAG]) or just the tRNA gene (pUltraI-Tet3.0 [TAG]) using the HiFi DNA assembly method (NEB #E2621L). In short, gBlocks containing at least 20 bp of homology to the pUltra backbone flanking genes for the previously reported wild-type *Methanosarcina mazei* pyrrolysyl-tRNA synthetase/tRNA pair (*Mm*PylRS/*Mm*tRNA_CUA_^Pyl^) recognizing the amber stop codon, were purchased from IDT. pUltra backbones were created using PCR amplification and purified using the Zymo DNA Clean and Concentrator PCR purification kit (Zymo #D4014). Fragments and backbones were ligated according to the NEBuilder HiFi DNA Assembly Master Mix protocol, transformed into chemically competent DH5α *E. coli* (NEB #C2987H), then plated at 37ºC on LB-Spec (50 mg/mL) selection media for 12-16 hours. Colonies were inoculated in LB-Spec (50 mg/mL) and grown at 37ºC (250 rpm) for 12-16 hours, and then miniprepped using the Qiagen QIAprep Spin Miniprep Kit (Qiagen #27104). The sequence of the pUltraI-*Mm*Pyl [TAG] and pUltraI-Tet3.0 [TAG] constructs was then confirmed using Genewiz Plasmid-EZ whole plasmid sequencing.

### pRESS Plasmid Construction

To create the pRESS plasmids, the parental pET28b plasmid was modified to include aaRS/tRNA genes from the pUltra system using the HiFi DNA assembly method (NEB #E2621L). gBlocks containing at least 20 bp of homology to each other and the pET28b backbone flanking genes for the wild-type *Methanosarcina mazei* pyrrolysyl-tRNA synthetase/tRNA pair (*Mm*PylRS/*Mm*tRNA_CUA_^Pyl^) recognizing the amber stop codon and under control of the T7 promoter/lacI operator system, were purchased from IDT. The T7p-lacI-MmPylRS-T7t gene also contained *PaqCI* cut sites for replacement with other aaRSs of interest using NEBridge® Golden Gate Assembly Kit protocol (NEB #E1601L). pET28b backbones were created via PCR amplification and purified using the Zymo DNA Clean and Concentrator PCR purification kit (Zymo #D4014). Fragments and backbones were ligated according to the NEBuilder HiFi DNA Assembly Master Mix protocol, transformed into chemically competent DH10β E. coli (NEB #C3019H), then plated at 37ºC on LB-Kan (50 mg/mL) selection media for 12-16 hours. Colonies were inoculated in LB-Kan (50 mg/mL) and grown at 37ºC (250 rpm) for 12-16 hours, and then miniprepped using the Qiagen QIAprep Spin Miniprep Kit (Qiagen #27104). The sequence of the pRESS-*Mm*Pyl construct was then confirmed using Genewiz Plasmid-EZ whole plasmid sequencing. To insert the genes for MBP-mCherry, bdSUMO nPD-L1, bdSUMO nCD16, bdSUMO nCTLA-4, and all associated variants into pRESS-*Mm*Pyl, an identical method for ordering and assembling plasmids via *BsaI* Golden Gate Assembly, as described in the “pET28b Plasmid Construction” section, was followed. Similarly, to create the pRESS-Tet3.0 variant, a gBlock containing the gene for the previously described *Methanosarcina barkeri-*derived pyrrolysyl-tRNA synthetase (*Mb*Tet3.0RS) with flanking *PaqCI* cut sites was ordered from IDT and assembled via a Golden Gate reaction. All constructs were confirmed using Genewiz Plasmid-EZ whole plasmid sequencing.

### Bacterial Culture and Expression Conditions

All cultures were grown on the 50 to 100 mL scale in sterile baffled 500 mL culture flasks in a New Brunswick Innova® 44R incubator shaker set to 37ºC and 250 rpm. Confirmed constructs described in the “pET28b, pUltra, and pRESS Plasmid Construction” sections were transformed into BL21(DE3) competent E. coli (in the case of MBP-mCherry) (NEB #C2527H) or SHuffle^®^ T7 Express Competent *E. coli* (in the case of nanobodies) (NEB #C3029J) and either plated at 37ºC on LB agar selection media containing the appropriate antibiotics for 12-16 hours or directly used to inoculate 5 mL of overnight starter cultures in LB media supplemented with the appropriate antibiotics. For plated constructs, overnight starter cultures in LB media were inoculated by scraping a swath of cells from each plate. Overnight starter cultures in LB media supplemented with the appropriate antibiotics were used to inoculate expression cultures at 2% (v/v) dilution. All cultures for protein expression were performed in non-auto-inducing Bosco Broth (BB) (As prepared by Bosco et al) supplemented with Kan (50 mg/mL) and/or Spec (50 mg/mL)^37^. All cultures were grown until an OD of 1.5-2 was reached, at which point cells were cooled, induced with a final concentration of 0.3 mM Isopropyl β-D-1-thiogalactopyranoside (IPTG) (Fisher #BP1755-1), and left to shake at 250 rpm and 16ºC for approximately 24 hours. Cultures for the expression of proteins containing Tet3.0 (GCE4All Research Center, Oregon State University, product #1003) were supplemented with 100 mM stock solutions prepared in fresh DMF to a final concentration of 1 mM prior to inoculation with overnight starters. Cultures for the expression of proteins containing Azk (Millipore Sigma #914088) were supplemented with 200 mM stock solutions prepared in fresh 1 M NaOH (H_2_O), to a final concentration of 2 mM, prior to inoculation with overnight starters.

### Bacterial Protein Purification

Proteins were expressed according to the culture conditions detailed in the “Bacterial Culture and Expression Conditions” section. After expression, cultures were harvested by pelleting at 6,800 x g for 12 minutes. The culture supernatant was removed, and pelleted cells were either used immediately or were stored frozen at - 20ºC until needed. For lysis, pelleted cultures were resuspended in 5 mL of ice-cold lysis buffer (PBS (ThermoFisher #10010023) supplemented with 10 mM imidazole, 5% v/v glycerol, and Roche cOmplete protease inhibitor cocktail EDTA free (Millipore Sigma #11836153001), pH 7.4) per gram of cell paste. These resuspended cells were then sonicated at 18% amplitude 5 seconds on 5 seconds off for a total run time of 20 minutes using a VWR homogenizer ultrasonic 50 watt. The resulting lysate was clarified by centrifugation at 16,000 x g for 35 minutes in a chilled centrifuge. The clarified lysate was combined with 1 mL of HisPur™ Ni-NTA Resin (ThermoFisher #88221) and incubated with agitation at 4ºC for at least 1 hour. The resin was then washed with 40-50 column volumes of wash buffer (PBS, supplemented with 25 mM imidazole, pH 7.4) and eluted with 2-3 column volumes of elution buffer (PBS supplemented with 250 mM imidazole, pH 7.4). The eluted protein solution was buffer-exchanged into PBS using a 10 kDa MWCO Amicon® Ultra Centrifugal Filter (Milipore Sigma #UFC901008) according to the manufacturer’s instructions. IMAC elutions were further purified using the Superdex 200 Increase 10/300 GL column (Cytvia #28990944) on the ÄKTA pure™ chromatography system (Cytvia #29018224).

### SDS Page and Densitometry

Samples were combined 1:1 with 2x Laemmli Sample Buffer (BioRad #1610737) (65.8 mM Tris-HCl (pH 6.8), 2.1% SDS, 26.3% (w/v) glycerol, and 0.01% bromophenol blue, 5% (v/v) β-mercaptoethanol, pH 6.8) before incubation at 95-100ºC for approximately 5 minutes. Samples were then loaded (volumes ranging from 3-18 µL) onto Any kD™ Mini-PROTEAN® TGX™ Precast Protein Gels (BioRad #4569036) gels and run at 200 V for approximately 30 minutes. SDS-PAGE gels were soaked in 50:50 mixture of staining/destaining solution (50% (v/v) methanol, 40% (v/v) water, 10% (v/v) acetic acid) and 1x NativePAGE™ Cathode Buffer Additive (ThermoFisher #BN2002) for a minimum of 30 minutes following microwaving. Gels were soaked and microwaved in destain solution (50% (v/v) methanol, 40% (v/v) water, 10% (v/v) acetic acid) to rapidly remove stain. When necessary, densitometry and colocalization measurements were made using the ImageJ software suite and JACoP plugins.

### Protein Mass Spectrometry

Following the purification methods described in the “Bacterial Protein Purification” section, 10-100 µL of protein underwent several rounds of microdialysis into 10 mM ammonium acetate (Millipore Sigma #372331) using 10 kDa MWCO Pierce™ Microdialysis Plates (ThermoFisher #88260). Protein concentrations ranged but were at least 5 µM for all samples. 5 µL of the sample was loaded into a glass capillary and placed on the emitter of the Q Exactive™ Plus Hybrid Quadrupole-Orbitrap™ Mass Spectrometer for nano-electrospray ionization mass spectrometry (ESI-MS). Samples were taken at a range of resolutions (17.5k to 280k) with an m/z range of 1000 to 6000. Raw files were evaluated and deconvoluted using UniDec.

### Bispecific Conjugation Reactions

Conjugation reactions between proteins were performed overnight using a 1:1:1 molar ratio of purified Tet3.0 containing protein, Azk containing protein, and TCO-PEG12-DBCO (BroadPharm #BP-22423) flexible linker dissolved in DMSO. The total volume of DMSO present in the reaction never exceeded 1%. Reactions were performed in the dark at room temperature in 1xPBS (pH 7.4) with agitation. Conjugates were purified using the Superdex 200 Increase 10/300 GL column (Cytvia #28990944) on the ÄKTA pure™ chromatography system (Cytvia #29018224).

## AUTHOR INFORMATION

### Author Contributions

Roman Adomanis: Wrote the original draft of the manuscript, generated all figures and graphics for the manuscript, edited, revised, and approved the final version of the manuscript. Nathan Phan: Supported data generation for the manuscript. Grant Walter: Supported data generation for the manuscript. Blaise Kimmel: Edited, revised, and approved the final version of the manuscript, and acquired funding to support the work.

## Acknowledgements

This work was supported in part by The Ohio State University Center for Cancer Engineering-Curing Cancer Through Research in Engineering and Sciences. B.R.K. acknowledges financial support from the Prostate Cancer Foundation Young Investigator Award. All figures created with Biorender.com.

