## Supplemental Table 1 for "Modular Nanobody Conjugates with Controlled Topology Using Genetically Encoded Non-canonical Amino Acids"

### Supplementary Information

**Table S1**: Expected and Observed Protein Molecular Weights.

| | Expected Mass (Da) | Observed Mass (Da) | Difference (Da) |
|---|---|---|---|
| **nPDL1 WT** | 13336.92 | 13060 | 276.92 |
| **nPDL1 Q1Azk** | 13450.03 | 13173 | 277.03 |
| **nPDL1 R45Azk** | 13421.98 | 13144 | 277.98 |
| **nPDL1 K75Azk** | 13449.99 | 13175 | 274.99 |
| **nPDL1 S116Azk** | 13491.09 | 13214 | 277.09 |
| **nCTLA4 WT** | 14400.73 | 14124 | 276.73 |
| **nCTLA4 Q1Tet3.0** | 14555.93 | 14350 | 205.93 |
| **nCTLA4 S123Tet3.0** | 14597.02 | 14320 | 277.02 |
| **nCD16AB E1Tet3.0** | 14436.98 | 14436 | 0.98 |
| **Molecule (1)** | 28915.31 | 28610 | 305.31 |
| **Molecule (5)** | 29034.26 | 28877 | 157.26 |
| **Molecule (6)** | 29075.35 | 28877 | 198.35 |
| **Molecule (11)** | 29034.22 | 28566 | 468.22 |
